# Evolutionary diversity of the control of the azole response by Tra1 across yeast species

**DOI:** 10.1101/2023.09.05.556458

**Authors:** Gabriela Marsiglio Nunes Librais, Yuwei Jiang, Iqra Razzaq, Christopher J. Brandl, Rebecca S. Shapiro, Patrick Lajoie

**Author notes:** Corresponding authors: Patrick Lajoie, PhD, Department of Anatomy and Cell Biology, The University of Western Ontario London, Ontario N6A 5C1 Canada, Rebecca S. Shapiro, PhD, Department of Cell and Molecular Biology, University of Guelph, Guelph, Ontario N1G 2W1 Canada.

## Abstract

Tra1 is an essential co-activator protein of the yeast SAGA and NuA4 acetyltransferase complexes that regulate gene expression through multiple mechanisms including the acetylation of histone proteins. Tra1 is a pseudokinase of the PIKK family characterized by a C-terminal PI3K domain with no known kinase activity. However, mutations of specific arginine residues to glutamine in the PI3K domains (an allele termed *tra1*_*Q3*_) result in reduced growth and increased sensitivity to multiple stresses. In the opportunistic fungal pathogen *Candida albicans*, the *tra1*_*Q3*_ allele reduces pathogenicity and increases sensitivity to the echinocandin antifungal drug caspofungin, which disrupts the fungal cell wall. Here, we found that loss of Tra1 function, in contrast to what is seen with caspofungin, increases tolerance to the azole class of antifungal drugs, which inhibits ergosterol synthesis. In *C. albicans, tra1*_*Q3*_ increases expression of genes linked to azole resistance, such as *ERG11* and *CDR1. CDR1* encodes a multidrug ABC transporter associated with efflux of multiple xenobiotics, including azoles. Consequently, cells carrying *tra1*_*Q3*_ show reduced intracellular accumulation of fluconazole. In contrast, a *tra1*_*Q3*_ *S. cerevisiae* strain displayed opposite phenotypes: decreased tolerance to azole, decreased expression of the efflux pump *PDR5* and increased intracellular accumulation of fluconazole. Therefore, our data provide evidence that Tra1 differentially regulates the antifungal response across yeast species.

## INTRODUCTION

Fungal infections represent a major public health concern with over a billion infections each year resulting in more than 1.5 million deaths (Bongomin *et al* 2017). Members of the *Candida* genus, including *Candida albicans*, are opportunistic pathogens that can cause a wide range of severe infections in susceptible populations, such as the elderly or immunocompromised individuals (Pappas *et al*. 2018). *Candida* species represent the leading cause of fungal infection-related deaths worldwide (Rokas 2022). Treatment of *Candida* infection (candidiasis) is unfortunately limited to four major classes of antifungal drugs: echinocandins, azoles, polyenes and flucytosines. Echinocandins, such as caspofungin, target the fungal cell wall by inhibiting the synthesis of the carbohydrate β-1,3-glucan (Balashov *et al*. 2006; Lee *et al*. 2012), while azoles, such as fluconazole and miconazole, inhibit the synthesis of ergosterol, thereby compromising the lipid composition of fungal membranes (Kuznetsov 2021). Azoles specifically inhibit the 14α-demethylase, encoded by the *C. albicans ERG11* gene, in the ergosterol synthesis pathway. Consequently, mutations in *ERG11* are one of the main sources of acquired azole resistance (Wu *et al*. 2017; Kaur and Nobile 2022).

Tra1 is an essential component of the SAGA and NuA4 complexes that regulate the acetylation of both histone and non-histone substrates (Grant *et al*. *et al*. 1999; Steunou *et al*. 2014; Downey 2021) in all eukaryotic cells, including fungi (Saleh *et al*. 1998; Grant *et al*. 1998; Allard *et al*. 1999). Tra1 is essential due to its function in NuA4. This is highlighted in the fission yeast *Schizosaccharomyces pombe* where the SAGA-incorporated Tra1 is dispensable for viability while conversely, the Nua4-localized Tra2 is essential (Helmlinger *et al*. 2011). Many components of the SAGA and NuA4 complexes regulate different aspects of the *C. albicans* antifungal response and pathogenicity (Laprade *et al*. 2002; Lu *et al*. 2008; Sellam *et al*. 2009; Askew *et al*. 2009; Chang *et al*. 2015; Shivarathri *et al*. 2019; Razzaq *et al*. 2021; Rashid *et al*. 2022). However, the functions of Tra1 in *C. albicans* remain poorly understood.

Tra1 is a member of the PIKK (phosphoinositide-3-kinase-related kinase) family, but unlike other family members such as Tor1, it does not possess any detectable kinase activity due to the lack of specific kinase motifs (McMahon *et al*. 1998; Helmlinger *et al*. 2011). Despite its pseudokinase nature, we have previously shown that mutation of key arginine residues to glutamine in the Tra1 PI3K domain, an allele termed *tra1*_*Q3*_, is associated with increased sensitivity to various stress conditions, including cell wall stress, protein misfolding and high temperature in the budding yeast *Saccharomyces cerevisiae (Berg et al. 2018). tra1*_*Q3*_ mutants also display increased sensitivity to acid stress, impaired growth in respiratory conditions and reduced chronological lifespan (Bari *et* al. 2022). Similar to their *S. cerevisiae* counterparts, *C. albicans tra1*_*Q3*_ mutants display increased sensitivity to cell wall stress induced by caspofungin, as well as reduced biofilm formation and pathogenicity (Razzaq *et al*. 2021). However, the impact of Tra1 on *C. albicans* resistance to other classes of antifungal drugs remains unknown. Hence, here we tested the effect of the *tra1*_*Q3*_ mutation to modulate sensitivity to other antifungal compounds.

## RESULTS AND DISCUSSION

### Loss of Tra1 function differentially impact azole resistance across yeast species

To test the effect of the *tra1*_*Q3*_ allele on the *C. albicans* antifungal resistance, wild-type cells and cells carrying the mutations were spotted on agar plates containing either the echinocandin caspofungin, the azoles miconazole and fluconazole, or amphotericin B (Figure 1A and B). While caspofungin disrupts the fungal cell wall by inhibiting the β-(1,3)-D-glucan synthase, both azoles and amphotericin B affect fungal membranes. Azoles inhibit ergosterol synthesis by inhibiting the cytochrome P450 enzyme 1,4 ɑ-demethylase (Hitchcock 1991; Wu *et al*. 2017). Polyenes, such as amphotericin B, disrupts membrane integrity by directly binding to ergosterol (Nett *et al*. 2010). As previously observed (Razzaq *et al*. 2021), two independently generated *tra1*_*Q3*_ mutants display increased sensitivity to caspofungin. Similarly, we observed increased sensitivity to amphotericin B. Unexpectedly, *tra1*_*Q3*_ cells showed increased tolerance to the azoles miconazole and fluconazole compared to wild-type. While both azoles and amphotericin B affect fungal membrane integrity, they do so via very distinct mechanisms; mechanisms for development of antifungal resistance to these drugs are also very different (Lee *et al*. 2020). Interestingly, deleting other SAGA complex components such as *SPT7, SPT8* and *ADA2* sensitize *C. albicans* to both caspofungin and fluconazole (Bruno *et al*. 2006; Sellam *et al*. 2009; Rashid *et al*. 2022), whereas deleting other components, such as *NGG1* and *UPB8*, have no effect on antifungal drug resistance (Rashid *et al*. 2022). Therefore, different SAGA components and/or modules may differentially contribute to antifungal drug resistance. Moreover, due to its incorporation in both SAGA and NuA4, Tra1 has additional roles that may contribute to the phenotypes observed here. Indeed, while the role for NuA4 in hyphal growth has been characterized (Lu *et al*. 2008), how it regulates antifungal drug resistance remains unclear.

**Figure 1:**
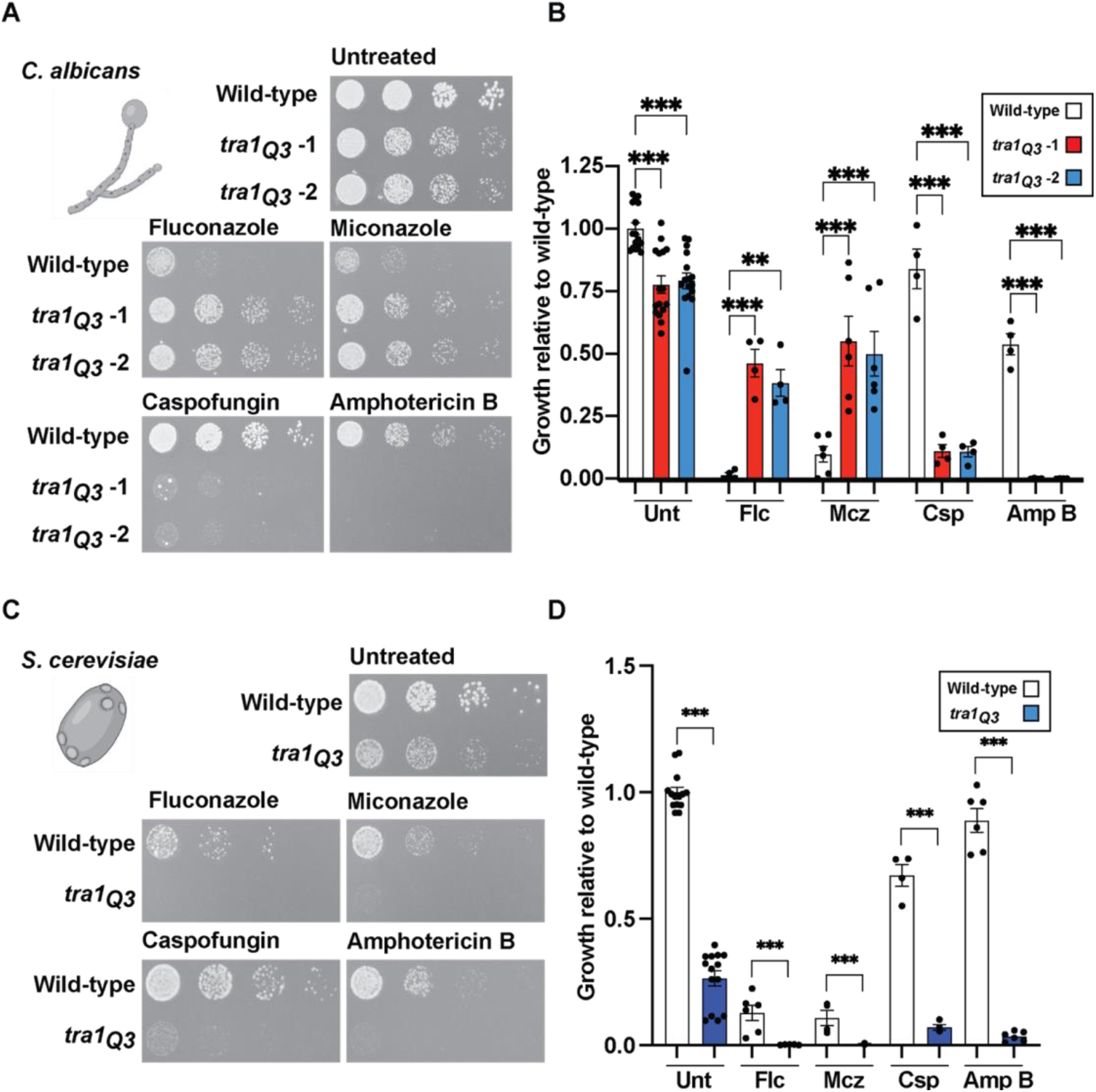
*tra1*_*Q3*_ results in differential resistance to azoles across yeast species. **(A)** Wild-type and *tra1*_*Q3*_ *C. albicans* cells were spotted on YPD agar plates at 30°C without treatment or containing either 20 µg/mL fluconazole (Flc), 0.5 µg/mL miconazole (Mcz), 0.05 µg/mL caspofungin (Csp) or 0.5 µg/mL amphotericin B (Amp B). **(B)** Quantification of the growth relative to untreated wild-type is shown in the bar graph. The second dilution was used for quantification. Data are presented ± sem n ≥ 4 and ***P<0.0003; **P<0.01 **(C)** Wild-type and *tra1*_*Q3*_ *S. cerevisiae* cells were spotted on YPD agar plates at 30°C without treatment or containing either 20 µg/mL fluconazole (Flc), 0.3 µg/mL miconazole (Mcz), 0.10 µg/mL caspofungin (Csp) or 0.5 µg/mL amphotericin B (Amp B). **(D)** Quantification of the growth relative to untreated wild-type is shown in the bar graph. Second dilution was used for quantification. Data are presented ± sem n ≥ 4 and ***P<0.0003; **P<0.01

In contrast, in *S. cerevisiae, tra1*_*Q3*_ cells are hypersensitive to caspofungin, amphotericin B and azoles (Figure 1C and D). This is consistent with previous results showing that loss of Tra1 function in budding yeast sensitizes cells to multiple stresses such as heat shock, protein misfolding, aging, cell wall perturbation and DNA damage (Mutiu *et al*. 2007; Hoke *et al*. 2008, 2010; Berg *et al*. 2018; Cheung and Díaz-Santín 2019; Jiang *et al*. 2019; Bari *et al*. 2022). The differential sensitivity to azoles between *C. albicans* and *S. cerevisiae* suggests that differences exist between the genes impacted by Tra1 across yeast species.

### Tra1 regulates the expression of genes associated with azole resistance

Given the decreased azole susceptibility in *tra1*_*Q3*_ cells, we next assessed the expression of genes previously associated with azole resistance in *C. albicans*. Specifically, we addressed the expression of *ERG11* and *CDR1*. Mutations in *ERG11* resulting in overexpression or loss of drug affinity are associated with azole resistance in multiple *C. albicans* clinical isolates (Franz *et al*. 1998; MacPherson *et al*. 2005; Flowers *et al*. 2015). Cdr1 is a member of the ATP-binding cassette transporter family associated with multidrug resistance (Dogra *et al*. 1999; Holmes *et al*. 2008). Cdr1 is known for its role in fluconazole efflux and acquired multidrug resistance in clinical isolates of *C. albicans (Holmes et al. 2008)*. As shown in Figure 2A, there was a significant increase in expression of *CDR1* and *ERG11* in *tra1*_*Q3*_ cells relative to wild-type both in untreated conditions (both 2.8 fold) and upon addition of fluconazole (2.7 and 2.1 fold respectively). In the past, we have shown, in budding yeast, that components of the SAGA complex can act as repressors of transcription under different conditions (Ricci *et al*. 2002).

**Figure 2:**
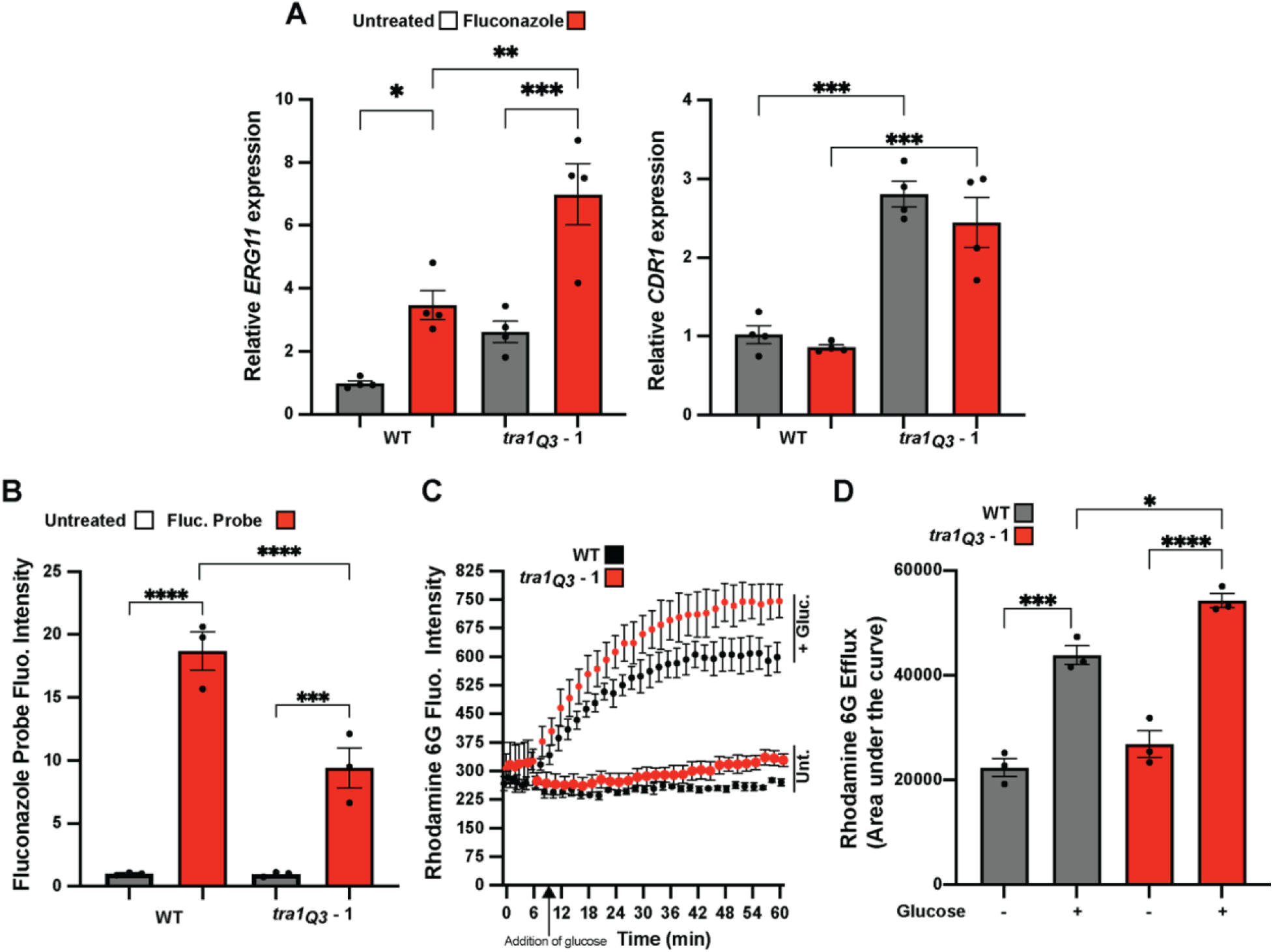
*C. albicans tra1*_*Q3*_ mutants display phenotypes associated with azole resistance. (**A**) *tra1*_*Q3*_ cells show increased expression of genes linked to azole resistance. Wild-type and *tra1*_*Q3*_ *C. albicans* cells were incubated with 20 µg/ml fluconazole for 1h and expression of *ERG11* and *CDR1* were assessed by qRT-PCR. n=4. (**B**) *tra1*_*Q3*_ cells show reduced accumulation of intracellular fluconazole. Wild-type and *tra1*_*Q3*_ *C. albicans* cells were incubated with a fluorescent fluconazole probe for 1h and mean fluorescent intensity of intracellular fluconazole was assessed by flow cytometry. n=3. (**C**) Rhodamine 6G efflux is increased in tra1_Q3_ cells. Energy-depleted cells were incubated with rhodamine 6G for 1h and then left untreated or treated with glucose to induce efflux. Mean rhodamine 6D fluorescence was monitored over time. n= 3 (**D**) Quantification of the area under the curve (AUC) calculated from rhodamine 6G release assays is shown in the bar graph. n=3. Means are shown ± sem ****P<0.0001; ***P<0.0003; **P<0.01; *P<0.05

In light of the increased expression of *CRD1* in *tra1*_*Q3*_ cells in *C. albicans*, we investigated whether the loss of Tra1 function affects drug efflux and intracellular bioavailability. To do so, we took advantage of a fluorescent probe that allows the real-time imaging of azole uptake in fungal cells (Benhamou *et al*. 2017). We found that intracellular accumulation of fluorescently-tagged azole is significantly reduced by 50% in *tra1*_*Q3*_ cells (Figure 2B). Since *CDR1* expression has been linked to increased efflux of fluconazole in fungi (Hernáez *et al*. 1998; Kim *et al*. 2019), we tested whether *tra1*_*Q3*_ *C. albicans* have higher drug efflux capacity using the well-characterized Cdr1 efflux substrate rhodamine 6G (Maesaki *et al*. 1999). Our findings support that there is increased efflux of rhodamine 6G in *tra1*_*Q3*_ cells (Figure 2C and D), suggesting that elevated expression of *CDR1* contributes to greater azole tolerance. This upregulation of efflux pumps may explain the unique azole-resistance of *tra1*_*Q3*_., Upregulation of efflux pumps are a well-established mechanism of azole resistance, but have not been linked to resistance of amphotericin B or caspofungin (Lee *et al*. 2020).

In response to azole, *C. albicans* activates the calcineurin pathway, which is essential for virulence (Blankenship *et al*. 2003; Juvvadi *et al*. 2014). Inhibiting calcineurin-dependent signaling with FK506 increases susceptibility to azole in *C. albicans* (Cruz et al. 2002; Uppuluri et al. 2008; *Khodavaisy et al. 2023). Thus, we tested whether FK506 alleviates the azole resistance observed in tra1*_*Q3*_ cells. Indeed, we found that azole tolerance in *tra1*_*Q3*_ cells is suppressed by treatment with the calcineurin inhibitor (Figure S1). Together, our findings suggest that while some cellular mechanisms associated with azole resistance, such as *ERG11* and *CDR1* expression, are increased in *tra1*_*Q3*_ cells, drug tolerance still requires a functional calcineurin pathway. This is in agreement with previous studies showing that expression *CDR1* and ergosterol biosynthesis genes are independent of calcineurin signaling (Cruz *et al*. 2002; Jia *et al*. 2016).

Next, we investigated the effect of *tra1*_*Q3*_ on azole resistance mechanisms in budding yeast given the differences between *C. albicans* and *S. cerevisiae* (Figure 1). In *S. cerevisiae*, SAGA regulates gene expression in response to changes in sterol content (Dewhurst-Maridor *et al*. 2017). NuA4 is also linked to sterol metabolism and *eaf1Δ* cells display increased accumulation of ergosterol esters (Pham *et al*. 2022). Unlike in *C. albicans, S. cerevisiae tra1*_*Q3*_ cells did not demonstrate significant changes in *ERG11* expression, as compared to the wild-type (Figure 3A). This difference with our *C. albicans* data is consistent with a potential rewiring of the role of Tra1 in the transcription of sterol genes between species. Since the *tra1*_*Q3*_ cells showed increased expression of *CDR1* in *C. albicans*, we investigated its impact on the expression of the *PDR5* ABC transporter (the *CDR1* homologue) in *S. cerevisiae*. We found reduced expression of *PDR5* of *tra1*_*Q3*_ cells (Figure 3A). This is consistent with the role of SAGA in the regulation of *PDR5* expression (Gao *et al*. 2004) and reduced global transcription of SAGA-regulated genes previously observed in the *tra1*_*Q3*_ mutant (Berg *et al*. 2018). Consistent with reduced *PDR5* expression, intracellular accumulation of fluorescently-labeled fluconazole increased in *tra1*_*Q3*_ cells (Figure 3B).

**Figure 3:**
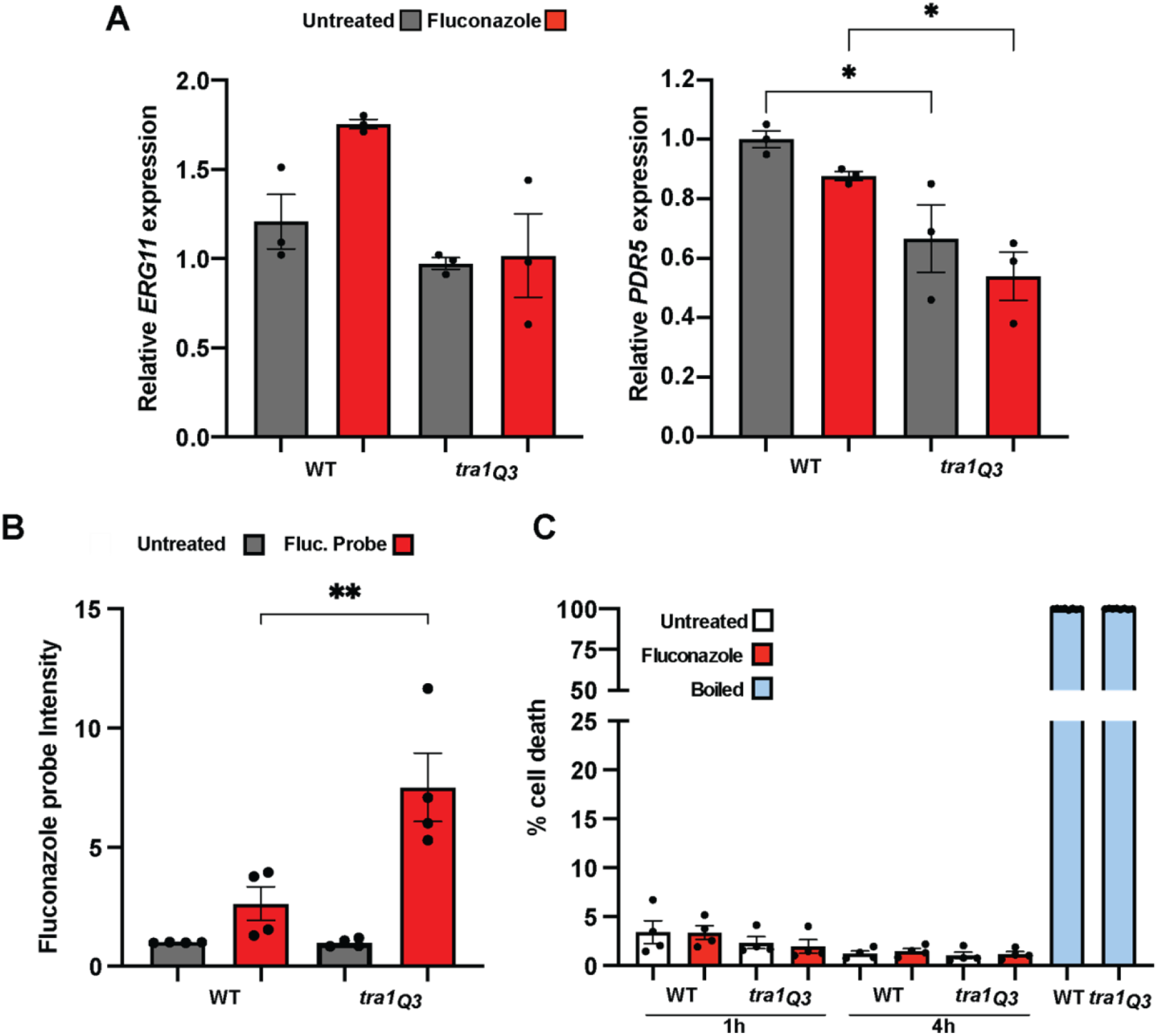
*S. cerevisiae tra1*_*Q3*_ mutants display phenotypes associated with increased azole sensitivity. (**A**) *tra1*_*Q3*_ cells show decreased expression of genes linked to azole resistance. Wild-type and *tra1*_*Q3*_ *C. albicans* cells were incubated with 20 µg/ml fluconazole for 1h and expression of *ERG11* and *PDR5* were assessed by qRT-PCR. n= 3. (**B**) *tra1*_*Q3*_ cells show increased accumulation of intracellular fluconazole. Wild-type and *tra1*_*Q3*_ cells were incubated with a fluorescent fluconazole probe for 1h and mean fluorescent intensity of intracellular fluconazole was assessed by flow cytometry. n=4. (**C**) *tra1*_*Q3*_ cells do not show increased cell death upon fluconazole treatment. Wild-type and *tra1*_*Q3*_ *S. cerevisiae* cells were incubated with 20 µg/ml fluconazole for 1 and 4 h and stained with propidium iodide (PI) to assess cell viability. Boiled cells are shown as positive control. The percentage of PI positive cells was assessed by flow cytometry and presented in a bar graph. n=4. All data are shown ± sem *P<0.05

**Figure 4:**
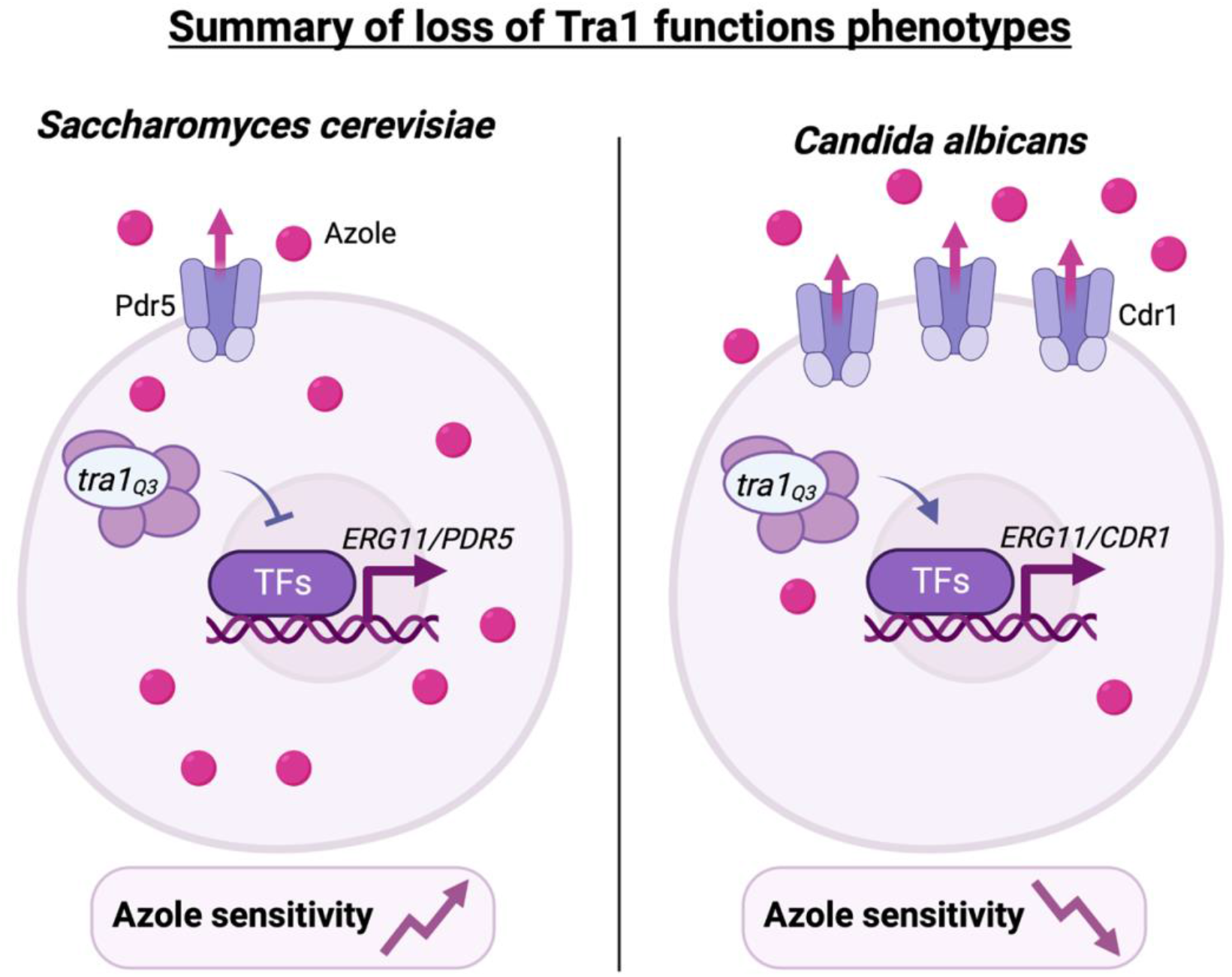
Summary of *tra1*_*Q3*_ phenotypes associated with azole treatment in *S. cerevisiae* and *C. albicans*. In *S. cerevisiae*, loss of Tra1 function associated with the *tra1*_*Q3*_ allele results in decreased expression of genes associated with azole resistance such as *ERG11* and *PDR5*. Consequently, *tra1*_*Q3*_ cells show increased accumulation of intracellular azole. In *C. albicans, tra1*_*Q3*_ is linked to increased expression of *ERG11* and *CDR1* and increased efflux of azole and consequently, increased drug resistance.

In contrast to other antifungal drugs, such as amphotericin B, which behave in a fungicidal manner, azoles are fungistatic, thus inducing minimal cell death in multiple yeast species (Manavathu *et al*. 1998). For this reason, we investigated whether the decreased growth observed in the *tra1*_*Q3*_ *S. cerevisiae* cells treated with fluconazole was associated with changes in cell death. Wild-type and *tra1*_*Q3*_ *S. cerevisiae* cells were treated with fluconazole and stained with PI at different time intervals for up to 24h (Figure 3C). No significant differences were observed, indicating that the effect of fluconazole remains fungistatic within the *tra1*_*Q3*_ cells. Finally, treatment of *S. cerevisiae* with the calcineurin inhibitor FK506 sensitized cells to fluconazole (Figure S2A). While *tra1*_*Q3*_ cells were not inherently sensitive to the inhibitor, the cells displayed a synthetic negative interaction when crossed with a *cnb1Δ* mutant (Figure S2B), which encodes a calcineurin subunit (Cyert and Thorner 1992). We previously observed a similar phenotype with a distinct Tra1 mutant (Hoke *et al*. 2008). These results suggest that Tra1 and calcineurin, like in *C. albicans*, function within distinct signaling pathways.

## Conclusions and perspectives

While biochemical and structural studies have extensively characterized Tra1 functions in model yeast, its specific impact for gene expression in fungal pathogens such as *C. albicans* is poorly understood. How loss of Tra1 function leads to upregulation of genes linked to azole resistance will require more detailed investigations to define genome-wide occupancy of coactivator complexes and their role in activation, repression and maintenance of transcription under different conditions.

Here, we also show that negatively impacting Tra1 function has opposite effects with regard to azole tolerance between *S. cerevisiae* and *C. albicans*. This highlights the evolutionary diversity of the control of the antifungal response by Tra1-containing complexes. Future investigations should aim at defining the breadth of Tra1 functions across fungi. *Nakaseomyces (Candida) glabrata* is the second most common cause of candidiasis (McCarty *et al*. 2021), but is evolutionarily more closely related to *S. cerevisiae* than other pathogenic *Candida* species. *N. glabrata* is highly dependent upon upregulation of *CDR1* and *CDR2* in response to antifungal stress (Sanglard *et al*. 2009), and thus could serve as a compelling comparison to assess this divergent role of Tra1 amongst yeast species. Finally, similar to other members of the PIKK family, such as Tor, Tra1 should be druggable. However, our data suggest that its potential as a candidate for combinational therapy with antifungal drugs would have to be considered carefully.

## MATERIAL AND METHODS

### Reagents

Fluconazole, amphotericin B, miconazole, FK506, rhodamine 6G and caspofungin were from MilliporeSigma. PI was from ThermoFisher Scientific.

### Yeast strains and growth conditions

The *tra1* mutant strains used in this study were previously described (Berg *et al*. 2018; Razzaq *et* al. 2021) and are listed in Table S1. Both *S. cerevisiae* and *C. albicans* strains were cultured in YPD (2% Bacto peptone, 1% yeast extract and 2% glucose) unless noted. Cells were grown in liquid YPD overnight at 30°C with shaking. The next day, cells were diluted 1:10 ratio and then incubated for 2 hr at 30°C with shaking. Cell growth on agar plates was measured as previously described (Petropavlovskiy *et al*. 2020).

### Fluorescent fluconazole probe uptake

The fluorescent fluconazole probe (RB510) was added to log phase cells cultured at at 30°C with shaking to a final concentration of 1µg/ml as previously described (Benhamou *et al*. 2017). Next, the cells were incubated in the dark for 60 min at 30°C with shaking. Then, cells were washed with phosphate-buffered saline (PBS). The mean fluorescence intensity of the probe was measured with a BD FacsCelesta flow cytometer. Data were collected from 30,000 cells per time point using a 561 nm yellow–green laser. Mean fluorescent intensity was calculated using Flowjo. No gates were applied.

### Cell viability assay

*S. cerevisiae* cells were grown in liquid SC media lacking leucine overnight at 30°C with shaking. The next day, cells were diluted at 1:5 ratio and then incubated for 5 hr at 30°C with shaking until the log phase in 50 mL culture flasks. Next, cells were equalized to OD_600_ = 0.80, treated or not with 20µg/ml of fluconazole and incubated with shaking for 1 to 24 hrs at 30°C. After each time point, 1 mL of cells were pelleted by centrifugation and resuspended in a final volume of 1000 µl PBS. For positive control, one sample of each strain was boiled for 15 min at 100°C. 500 µl of cell suspension were stained with 2.5µL of 1 mg/mL propidium PI solution, and cells were incubated in the dark for 10 min at room temperature as previously described (Chadwick *et al*. 2016). Data were collected from 30,000 cells per time point using a BD FACSCelesta flow cytometer (BD Biosciences) equipped with a 561 nm yellow-green laser. Analysis was performed using Flowjo.

### Gene expression analysis

Total RNA was isolated using the MasterPure™ Yeast RNA purification kit (Lucigen) according to the manufacturer’s instructions. cDNA was synthesized from 2.5 µg total RNA using the SuperScript™ IV VILO Master Mix (ThermoFisher Scientific) according to the manufacturer’s instructions. qRT-PCR for *ERG11* and *CDR1/PDR5* together with *ACT1/TDH3* as housekeeping gene were amplified from the synthesized cDNA using primers listed in Table S2 with a QuantStudio 3 real-time PCR system using the ΔΔCT method (ThermoFisher Scientific).

### Efflux of rhodamine 6G

To measure *C. albicans* drug efflux capacity, rhodamine 6G (R6G) efflux was measured by fluorescence assay with whole cells. *C. albicans* were grown in liquid YPD overnight at 30°C with shaking in 10 mL culture tubes. First, cells were diluted at a 1:10 ratio and then incubated for 2 hr at 30°C with shaking until the log phase. Next, cells were pelleted by centrifugation, washed with 5 ml PBS (pH 7), resuspended in 2 ml PBS, and incubated for 1 hr at 30°C with shaking in PBS to energy-deprived cells. R6G was added at a concentration of 4 µM, and the incubation continued for 1 hr, thus facilitating R6G accumulation. After this incubation, cells were sedimented by centrifugation, washed with PBS, and resuspended in a final volume of 200 µl PBS. Fifty microliters of individual strains were diluted in 150 µl PBS and aliquoted in a 96-well microtiter plate, which was placed in a BioTek Cytation5 Cell Imaging Multimode Reader (Agilent) with temperature control set at 30°C. Baseline emission of fluorescence (excitation wavelength: 584 nm; emission wavelength: 625 nm) and OD_600_ was recorded for 0, 2, and 4 min. Glucose (2% final concentration) was next added to each strain to initiate R6G efflux. As a negative control, no glucose was added to separate aliquots of each strain. Data points were recorded in triplicate for 60 min at 2-min intervals. Data were plotted as the ratio of fluorescence value/OD_600_ data point.

## DATA AVAILABILITY STATEMENT

Strains and plasmids are available upon request. The authors affirm that all data necessary for confirming the conclusions of the article are present within the article, figures, and tables.

## ACKNOWLEDGEMENTS

This study was supported by a Canadian Institutes for Health Research (CIHR) Project Grant (PJT 168882), a National Sciences and Engineering Research Council (NSERC) Discovery Grant (RGPIN-2022-05267) and a John R Evans Leader Fund Grant (65183) to PL. RSS is supported by a CIHR Project Grant (PJT 162195) and a NSERC Discovery Grant (RGPIN-2018-4914). CJB was supported by a NSERC Discovery Grant (RGPIN-2015-04394). YJ was supported by a Masters to PhD Transfer Scholarship from the Dean of the Schulich School of Medicine & Dentistry at Western University. The authors thank Micha Fridman (University of Tel-Aviv) for the fluorescent fluconazole probe. We also thank Matthew Berg for help in the early stages of the project.

## FIGURES

**Figure S1:**
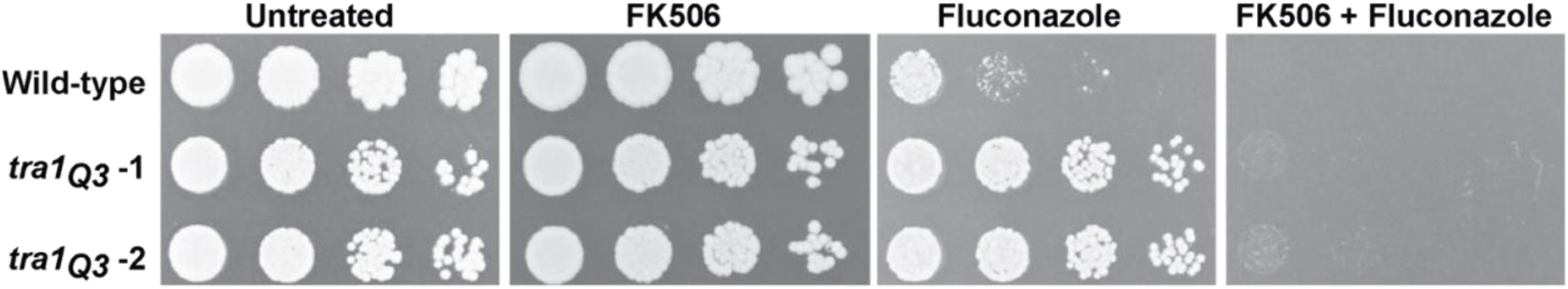
Treatment with FK506 alleviates the fluconazole resistance of *C. albicans tra1*_*Q3*_ cells. Wild-type and *tra1*_*Q3*_ *C. albicans* cells were spotted on YPD agar plates without treatment or containing either 20 µg/mL fluconazole, 2 µg/ml FK506 or a combination of both.

**Figure S2:**
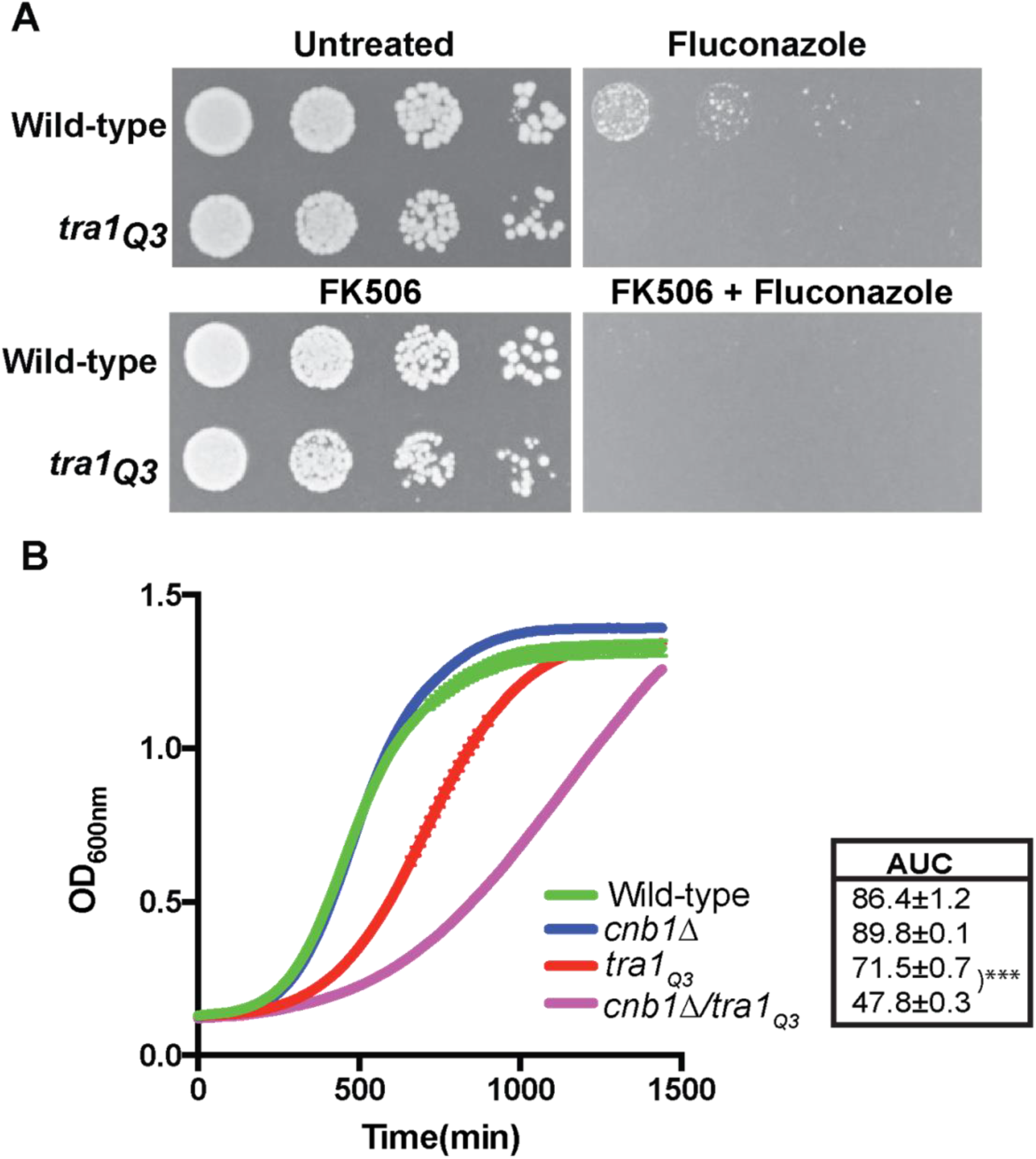
*S. cerevisiae tra1*_*Q3*_ shows synthetic negative genetic interaction with *CNB1*. **(A)** Wild-type, *cnb1Δ, tra1*_*Q3*_ and the double mutant *tra1*_*Q3*_ were spotted on YPD agar plates without treatment or containing either 20 µg/mL fluconazole, 2 µg/ml FK506 or a combination of both. **(B)** Wild-type, *cnb1Δ, tra1*_*Q3*_ and the double mutant were grown in YPD and OD_600_ was measured over time and growth curves were generated. For each strain the area under the curve was calculated and shown in insert.

**Table S1:**
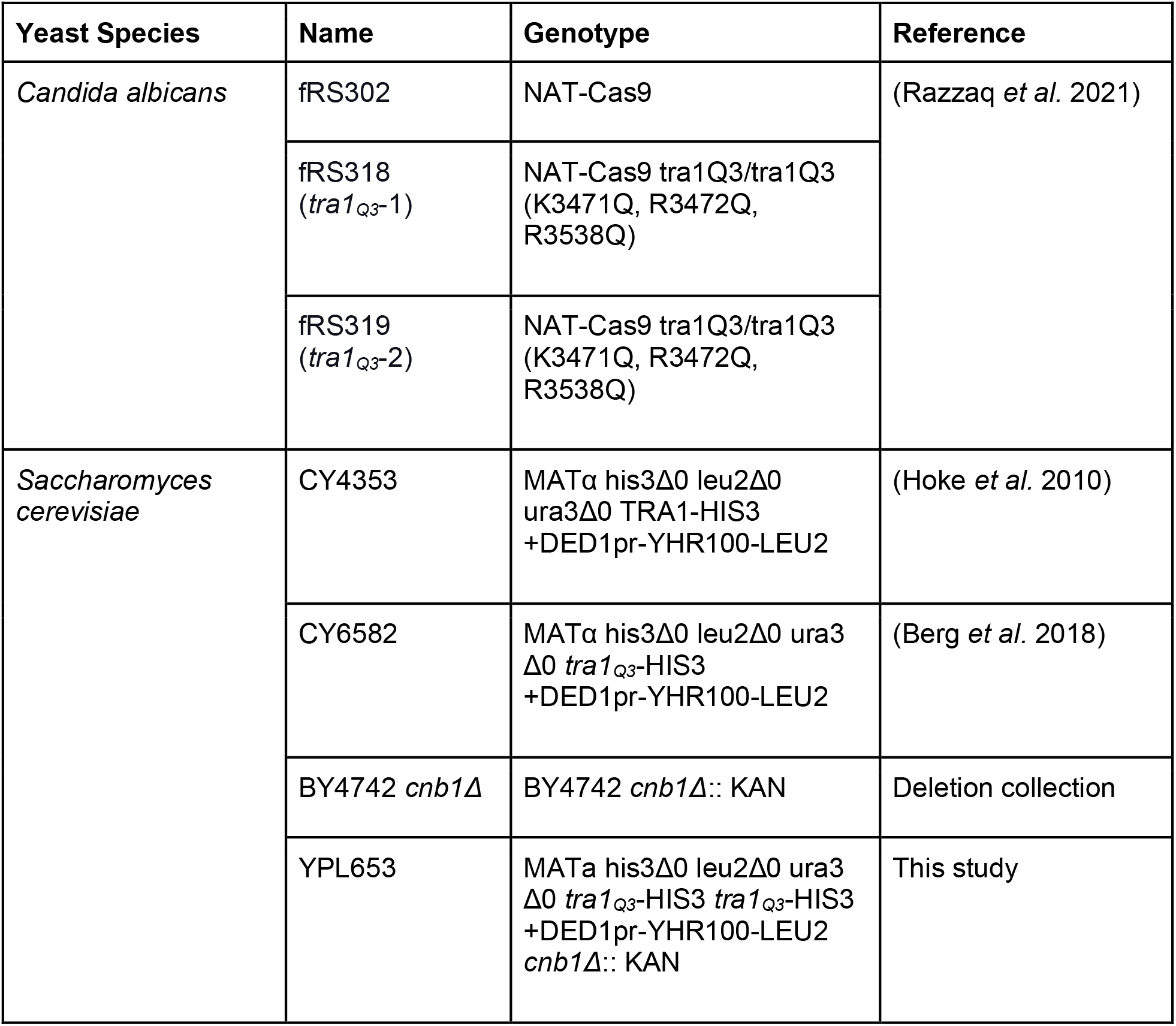
Strains used in this study.

**Table S2:**
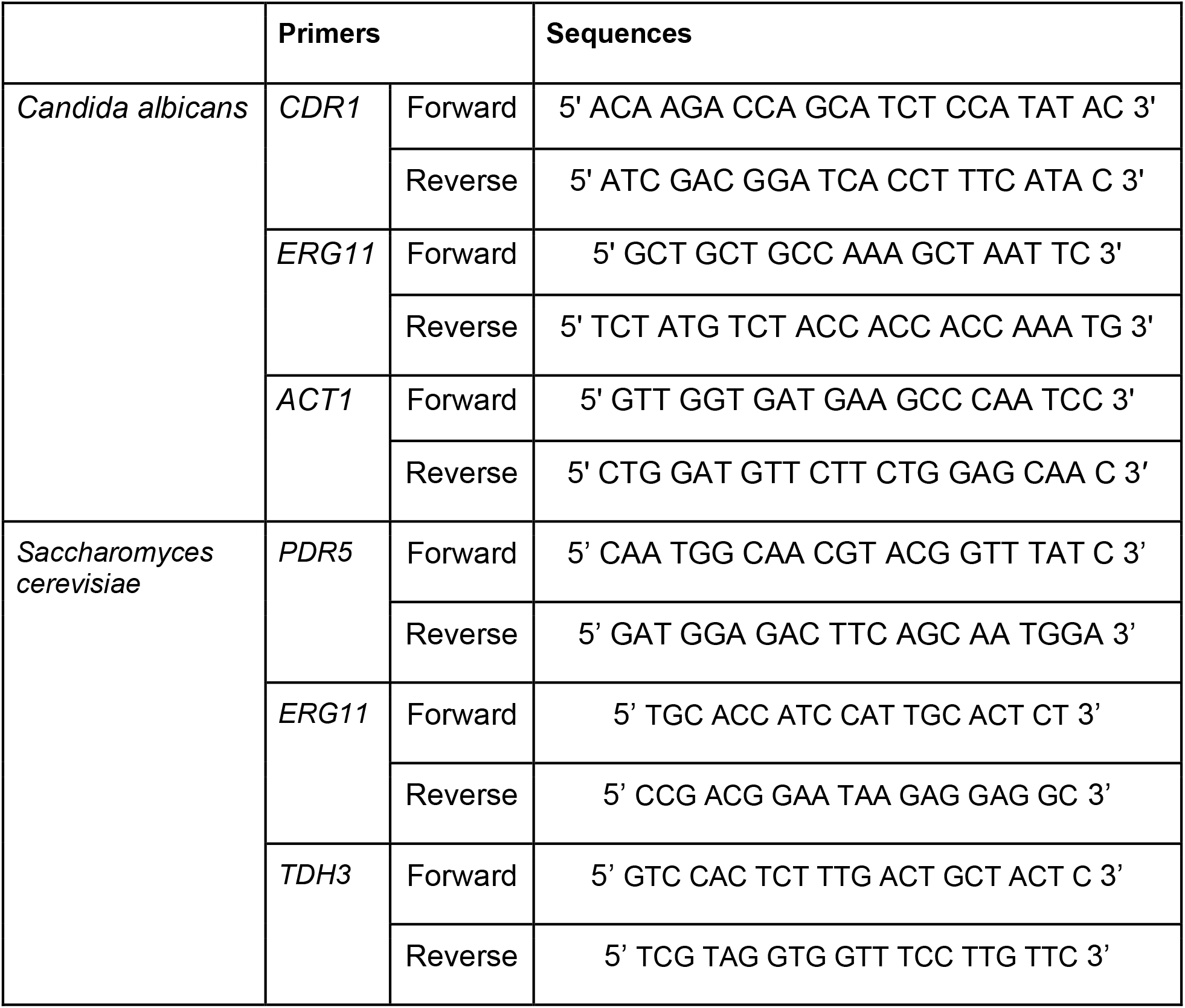
Primers used in this study.

## REFERENCES

Allard S., R. T. Utley, J. Savard, A. Clarke, P. Grant, et al., 1999 NuA4, an essential transcription adaptor/histone H4 acetyltransferase complex containing Esa1p and the ATM-related cofactor Tra1p. EMBO J. 18: 5108–5119.

Askew C., A. Sellam, E. Epp, H. Hogues, A. Mullick, et al., 2009 Transcriptional regulation of carbohydrate metabolism in the human pathogen Candida albicans. PLoS Pathog. 5: e1000612.

Balashov S. V., S. Park, and D. S. Perlin, 2006 Assessing resistance to the echinocandin antifungal drug caspofungin in Candida albicans by profiling mutations in FKS1. Antimicrob. Agents Chemother. 50: 2058–2063.

Bari K. A., M. D. Berg, J. Genereaux, C. J. Brandl, and P. Lajoie, 2022 Tra1 controls the transcriptional landscape of the aging cell. G3. 10.1093/g3journal/jkac287

Benhamou R. I., M. Bibi, K. B. Steinbuch, H. Engel, M. Levin, et al., 2017 Real-Time Imaging of the Azole Class of Antifungal Drugs in Live Candida Cells. ACS Chem. Biol. 12: 1769–1777.

Berg M. D., J. Genereaux, J. Karagiannis, and C. J. Brandl, 2018 The Pseudokinase Domain of Tra1 Is Required for Nuclear Localization and Incorporation into the SAGA and NuA4 Complexes. G3 8: 1943–1957.

Blankenship J. R., F. L. Wormley, M. K. Boyce, W. A. Schell, S. G. Filler, et al., 2003 Calcineurin is essential for Candida albicans survival in serum and virulence. Eukaryot. Cell 2: 422–430.

Bongomin F., S. Gago, R. O. Oladele, and D. W. Denning, 2017 Global and Multi-National Prevalence of Fungal Diseases-Estimate Precision. J Fungi (Basel) 3. 10.3390/jof3040057

Bruno V. M., S. Kalachikov, R. Subaran, C. J. Nobile, C. Kyratsous, et al., 2006 Control of the C. albicans cell wall damage response by transcriptional regulator Cas5. PLoS Pathog. 2: e21.

Chadwick S. R., A. D. Pananos, S. E. Di Gregorio, A. E. Park, P. Etedali-Zadeh, et al., 2016 A Toolbox for Rapid Quantitative Assessment of Chronological Lifespan and Survival in Saccharomyces cerevisiae. Traffic 17: 689–703.

Chang P., X. Fan, and J. Chen, 2015 Function and subcellular localization of Gcn5, a histone acetyltransferase in Candida albicans. Fungal Genet. Biol. 81: 132–141.

Cheung A. C. M., and L. M. Díaz-Santín, 2019 Share and share alike: the role of Tra1 from the SAGA and NuA4 coactivator complexes. Transcription 10: 37–43.

Clarke A. S., J. E. Lowell, S. J. Jacobson, and L. Pillus, 1999 Esa1p is an essential histone acetyltransferase required for cell cycle progression. Mol. Cell. Biol. 19: 2515–2526.

Cruz M. C., A. L. Goldstein, J. R. Blankenship, M. Del Poeta, D. Davis, et al., 2002 Calcineurin is essential for survival during membrane stress in Candida albicans. EMBO J. 21: 546–559.

Cyert M. S., and J. Thorner, 1992 Regulatory subunit (CNB1 gene product) of yeast Ca2+/calmodulin-dependent phosphoprotein phosphatases is required for adaptation to pheromone. Mol. Cell. Biol. 12: 3460–3469.

Dewhurst-Maridor G., D. Abegg, F. P. A. David, J. Rougemont, C. C. Scott, et al., 2017 The SAGA complex, together with transcription factors and the endocytic protein Rvs167p, coordinates the reprofiling of gene expression in response to changes in sterol composition in. Mol. Biol. Cell 28: 2637–2649.

Dogra S., S. Krishnamurthy, V. Gupta, B. L. Dixit, C. M. Gupta, et al., 1999 Asymmetric distribution of phosphatidylethanolamine in C. albicans: possible mediation by CDR1, a multidrug transporter belonging to ATP binding cassette (ABC) superfamily. Yeast 15: 111–121.

Downey M., 2021 Non-histone protein acetylation by the evolutionarily conserved GCN5 and PCAF acetyltransferases. Biochim. Biophys. Acta Gene Regul. Mech. 1864: 194608.

Flowers S. A., B. Colón, S. G. Whaley, M. A. Schuler, and P. D. Rogers, 2015 Contribution of clinically derived mutations in ERG11 to azole resistance in Candida albicans. Antimicrob. Agents Chemother. 59: 450–460.

Franz R., S. L. Kelly, D. C. Lamb, D. E. Kelly, M. Ruhnke, et al., 1998 Multiple molecular mechanisms contribute to a stepwise development of fluconazole resistance in clinical Candida albicans strains. Antimicrob. Agents Chemother. 42: 3065–3072.

Gao C., L. Wang, E. Milgrom, and W.-C. W. Shen, 2004 On the mechanism of constitutive Pdr1 activator-mediated PDR5 transcription in Saccharomyces cerevisiae: evidence for enhanced recruitment of coactivators and altered nucleosome structures. J. Biol. Chem. 279: 42677–42686.

Grant P. A., L. Duggan, J. Côté, S. M. Roberts, J. E. Brownell, et al., 1997 Yeast Gcn5 functions in two multisubunit complexes to acetylate nucleosomal histones: characterization of an Ada complex and the SAGA (Spt/Ada) complex. Genes Dev. 11: 1640–1650.

Grant P. A., D. Schieltz, M. G. Pray-Grant, J. R. Yates 3rd, and J. L. Workman, 1998 The ATM-related cofactor Tra1 is a component of the purified SAGA complex. Mol. Cell 2: 863–867.

Helmlinger D., S. Marguerat, J. Villén, D. L. Swaney, S. P. Gygi, et al., 2011 Tra1 has specific regulatory roles, rather than global functions, within the SAGA co-activator complex. EMBO J. 30: 2843–2852.

Hernáez M. L., C. Gil, J. Pla, and C. Nombela, 1998 Induced expression of the Candida albicans multidrug resistance gene CDR1 in response to fluconazole and other antifungals. Yeast 14: 517–526.

Hitchcock C. A., 1991 Cytochrome P-450-dependent 14 alpha-sterol demethylase of Candida albicans and its interaction with azole antifungals. Biochem. Soc. Trans. 19: 782–787.

Hoke S. M. T., J. Guzzo, B. Andrews, and C. J. Brandl, 2008 Systematic genetic array analysis links the Saccharomyces cerevisiae SAGA/SLIK and NuA4 component Tra1 to multiple cellular processes. BMC Genet. 9: 46.

Hoke S. M. T., A. Irina Mutiu, J. Genereaux, S. Kvas, M. Buck, et al., 2010 Mutational analysis of the C-terminal FATC domain of Saccharomyces cerevisiae Tra1. Curr. Genet. 56: 447–465.

Holmes A. R., Y.-H. Lin, K. Niimi, E. Lamping, M. Keniya, et al., 2008 ABC transporter Cdr1p contributes more than Cdr2p does to fluconazole efflux in fluconazole-resistant Candida albicans clinical isolates. Antimicrob. Agents Chemother. 52: 3851–3862.

Jia W., H. Zhang, C. Li, G. Li, X. Liu, et al., 2016 The calcineruin inhibitor cyclosporine a synergistically enhances the susceptibility of Candida albicans biofilms to fluconazole by multiple mechanisms. BMC Microbiol. 16: 113.

Jiang Y., M. D. Berg, J. Genereaux, K. Ahmed, M. L. Duennwald, et al., 2019 Sfp1 links TORC1 and cell growth regulation to the yeast SAGA-complex component Tra1 in response to polyQ proteotoxicity. Traffic 20: 267–283.

Juvvadi P. R., F. Lamoth, and W. J. Steinbach, 2014 Calcineurin as a Multifunctional Regulator: Unraveling Novel Functions in Fungal Stress Responses, Hyphal Growth, Drug Resistance, and Pathogenesis. Fungal Biol. Rev. 28: 56–69.

Kaur J., and C. J. Nobile, 2022 Antifungal drug-resistance mechanisms in Candida biofilms. Curr. Opin. Microbiol. 71: 102237.

Khodavaisy S., S. A. Gharehbolagh, M. Abdorahimi, S. Rezaie, K. Ahmadikia, et al., 2023 In vitro combination of antifungal drugs with tacrolimus (FK506) holds promise against clinical Candida species, including Candida auris. Med. Mycol. 61. 10.1093/mmy/myad069

Kim S. H., K. R. Iyer, L. Pardeshi, J. F. Muñoz, N. Robbins, et al., 2019 Genetic Analysis of Implicates Hsp90 in Morphogenesis and Azole Tolerance and Cdr1 in Azole Resistance. MBio 10. 10.1128/mBio.02529-18

Kuznetsov A. E., 2021 Introductory Chapter: Azoles, Their Importance, and Applications. Azoles - Synthesis, Properties, Applications and Perspectives.

Laprade L., V. L. Boyartchuk, W. F. Dietrich, and F. Winston, 2002 Spt3 plays opposite roles in filamentous growth in Saccharomyces cerevisiae and Candida albicans and is required for C. albicans virulence. Genetics 161: 509–519.

Lee K. K., D. M. Maccallum, M. D. Jacobsen, L. A. Walker, F. C. Odds, et al., 2012 Elevated cell wall chitin in Candida albicans confers echinocandin resistance in vivo. Antimicrob. Agents Chemother. 56: 208–217.

Lee Y., E. Puumala, N. Robbins, and L. E. Cowen, 2020 Antifungal Drug Resistance: Molecular Mechanisms in Candida albicans and Beyond. Chem. Rev. 10.1021/acs.chemrev.0c00199

Lu Y., C. Su, X. Mao, P. P. Raniga, H. Liu, et al., 2008 Efg1-mediated recruitment of NuA4 to promoters is required for hypha-specific Swi/Snf binding and activation in Candida albicans. Mol. Biol. Cell 19: 4260–4272.

MacPherson S., B. Akache, S. Weber, X. De Deken, M. Raymond, et al., 2005 Candida albicans Zinc Cluster Protein Upc2p Confers Resistance to Antifungal Drugs and Is an Activator of Ergosterol Biosynthetic Genes. Antimicrobial Agents and Chemotherapy 49: 1745–1752.

Maesaki S., P. Marichal, H. Vanden Bossche, D. Sanglard, and S. Kohno, 1999 Rhodamine 6G efflux for the detection of CDR1-overexpressing azole-resistant Candida albicans strains. J. Antimicrob. Chemother. 44: 27–31.

Manavathu E. K., J. L. Cutright, and P. H. Chandrasekar, 1998 Organism-dependent fungicidal activities of azoles. Antimicrob. Agents Chemother. 42: 3018–3021.

McCarty T. P., C. M. White, and P. G. Pappas, 2021 Candidemia and Invasive Candidiasis. Infect. Dis. Clin. North Am. 35: 389–413.

McMahon S. B., H. A. Van Buskirk, K. A. Dugan, T. D. Copeland, and M. D. Cole, 1998 The novel ATM-related protein TRRAP is an essential cofactor for the c-Myc and E2F oncoproteins. Cell 94: 363–374.

Mutiu A. I., S. M. T. Hoke, J. Genereaux, C. Hannam, K. MacKenzie, et al., 2007 Structure/function analysis of the phosphatidylinositol-3-kinase domain of yeast tra1. Genetics 177: 151–166.

Nett J. E., K. Crawford, K. Marchillo, and D. R. Andes, 2010 Role of Fks1p and matrix glucan in Candida albicans biofilm resistance to an echinocandin, pyrimidine, and polyene. Antimicrob. Agents Chemother. 54: 3505–3508.

Pappas P. G., M. S. Lionakis, M. C. Arendrup, L. Ostrosky-Zeichner, and B. J. Kullberg, 2018 Invasive candidiasis. Nat Rev Dis Primers 4: 18026.

Petropavlovskiy A. A., M. G. Tauro, P. Lajoie, and M. L. Duennwald, 2020 A Quantitative Imaging-Based Protocol for Yeast Growth and Survival on Agar Plates. STAR Protoc 1: 100182.

Pham T., E. Walden, S. Huard, J. Pezacki, M. D. Fullerton, et al., 2022 Fine-tuning acetyl-CoA carboxylase 1 activity through localization: functional genomics reveals a role for the lysine acetyltransferase NuA4 and sphingolipid metabolism in regulating Acc1 activity and localization. Genetics 221. 10.1093/genetics/iyac086

Rashid S., T. O. Correia-Mesquita, P. Godoy, R. P. Omran, and M. Whiteway, 2022 SAGA Complex Subunits in Differentially Regulate Filamentation, Invasiveness, and Biofilm Formation. Front. Cell. Infect. Microbiol. 12: 764711.

Razzaq I., M. D. Berg, Y. Jiang, J. Genereaux, D. Uthayakumar, et al., 2021 The SAGA and NuA4 component Tra1 regulates Candida albicans drug resistance and pathogenesis. Genetics 219. 10.1093/genetics/iyab131

Ricci A. R., J. Genereaux, and C. J. Brandl, 2002 Components of the SAGA histone acetyltransferase complex are required for repressed transcription of ARG1 in rich medium. Mol. Cell. Biol. 22: 4033–4042.

Rokas A., 2022 Evolution of the human pathogenic lifestyle in fungi. Nat Microbiol 7: 607–619.

Saleh A., D. Schieltz, N. Ting, S. B. McMahon, D. W. Litchfield, et al., 1998 Tra1p is a component of the yeast Ada.Spt transcriptional regulatory complexes. J. Biol. Chem. 273: 26559–26565.

Sanglard D., A. Coste, and S. Ferrari, 2009 Antifungal drug resistance mechanisms in fungal pathogens from the perspective of transcriptional gene regulation. FEMS Yeast Res. 9: 1029–1050.

Sellam A., C. Askew, E. Epp, H. Lavoie, M. Whiteway, et al., 2009 Genome-wide mapping of the coactivator Ada2p yields insight into the functional roles of SAGA/ADA complex in Candida albicans. Mol. Biol. Cell 20: 2389–2400.

Shivarathri R., M. Tscherner, F. Zwolanek, N. K. Singh, N. Chauhan, et al., 2019 The Fungal Histone Acetyl Transferase Gcn5 Controls Virulence of the Human Pathogen Candida albicans through Multiple Pathways. Sci. Rep. 9: 9445.

Steunou A.-L., D. Rossetto, and J. Côté, 2014 Regulating Chromatin by Histone Acetylation. Fundamentals of Chromatin 147–212.

Uppuluri P., J. Nett, J. Heitman, and D. Andes, 2008 Synergistic effect of calcineurin inhibitors and fluconazole against Candida albicans biofilms. Antimicrob. Agents Chemother. 52: 1127–1132.

Wu Y., N. Gao, C. Li, J. Gao, and C. Ying, 2017 A newly identified amino acid substitution T123I in the 14α-demethylase (Erg11p) of Candida albicans confers azole resistance. FEMS Yeast Res. 17. 10.1093/femsyr/fox012

